# A Population Coupling Model Identifies Reduced Propagation from V1 to Higher Visual Areas During Locomotion

**DOI:** 10.64898/2026.02.04.703681

**Authors:** Qi Xin, Konrad N. Urban, Joshua H. Siegle, Robert E. Kass

## Abstract

Point process generalized linear models (GLMs) have been a major tool for studying coordinated activity across populations of neurons. These models typically quantify how the spiking of a single neuron depends on the past activity of other neurons at multiple time lags, and the resulting neuron-to-neuron interactions are then aggregated to obtain population coupling effects. However, when neurons within the same population exhibit similar spiking patterns, explicitly modeling individual interactions can become redundant and unnecessarily increase model complexity. In such cases, population-level formulations may offer a more efficient alternative. For example, biophysical population models often characterize circuit dynamics using the average firing rate across neurons within a population, and recent data-driven approaches have similarly demonstrated the utility of population-level statistics for capturing cross-population interactions. Motivated by this consideration, we reformulate the GLM framework to operate directly at the population level. The resulting model, which we call pop-GLM, provides a computationally efficient method for estimating coupling between populations. In a simulated dataset, we show that pop-GLM achieves greater sensitivity in detecting coupling effects and can account for trial-to-trial variation in stimulus drive, which would otherwise introduce bias. We introduce a modification to the conventional GLM structure that is required to achieve the population-level formulation. We then apply pop-GLM to real data and identify decreased functional connectivity from primary visual area (V1) to a higher visual area during locomotion, a change not detected by single-neuron GLMs.

**Author summary:** A central goal of systems neuroscience is to understand how multiple populations of neurons, across different brain areas, interact as a coordinated circuit to produce perception and behavior. A common way to measure such interactions is to model how each neuron’s activity depends on the recent activity of others, and then combine these neuron-by-neuron estimates. When neurons in an area respond similarly, however, this becomes inefficient and hard to interpret. We instead formulated and investigated a new method that estimates functional interactions directly between two whole populations of spiking neurons, and showed it can be more sensitive and robust than previous approaches. To illustrate, we discovered decreased interaction between two mouse visual areas during locomotion, a change that previous techniques could not detect. The method should aid investigators in searching for important functional relationships across populations of neurons, with precise time-scale resolution.

## 1 Introduction

As simultaneous recording of neural activity across large numbers of neurons and brain regions becomes increasingly common [1; 2; 3; 4], interest has grown in modeling interactions at the population level to understand coordination across circuits [5; 6; 7]. A widely used statistical approach for studying such interactions is the point process generalized linear model (GLM), which models how a single neuron’s spiking is influenced by the past activity of other neurons, typically across multiple time lags [5; 8]. To characterize interactions between populations, it is common to aggregate the resulting neuron-to-neuron couplings across populations [9; 10; 11; 12].

In sensory cortical areas, subsets of neurons can often be identified that exhibit similar stimulus-evoked spiking patterns, allowing them to be treated as functionally defined populations. When this is the case, distinguishing between individual neurons is typically not the goal, since their firing tends to reflect a common underlying pattern; instead, the focus shifts to capturing interactions at the population level, which are more informative for understanding circuit computation. Moreover, modeling each neuron separately introduces substantial complexity: the number of coupling parameters grows quadratically with the number of neurons in the population, leading to noisy estimates that are difficult to interpret. Consistent with this perspective, population-level biophysical models have long focused on average firing rate within each population and have been successful in explaining various circuit-level phenomena [13; 14; 15; 16]. These models motivate a complementary statistical perspective: by defining populations as groups of neurons with similar spiking patterns, the most critical interactions may be better captured directly at the population level. Supporting this view, recent data-driven studies showed that the peak timing of population firing rate reveals strong temporal correlations and precise time lag estimates across populations [17; 18], suggesting that population-level statistics can meaningfully capture cross-population coupling.

Motivated by these developments, we present a population-level GLM coupling model (pop-GLM) that directly estimates interactions between populations using population spike trains obtained by pooling spike trains across neurons. Although this approach sacrifices individual neuron identity and requires grouping neurons with similar firing patterns, pop-GLM avoids the need to estimate a large number of neuron-to-neuron couplings and is therefore more sensitive to detecting coupling effects. In addition, pooled spike trains contain richer information than individual spike trains and allow us to estimate and adjust for trial-to-trial variability arising from non-coupling sources, such as locomotion or attention. Beyond the methodological contribution, we apply pop-GLM to large-scale recordings from mouse visual cortex and find that locomotion is associated with decreased functional connectivity between visual areas. This observation complements existing findings of locomotion-induced modulation of firing rate and response variability [19; 20; 21; 22; 23; 24; 25; 26; 27; 28; 29; 30], providing new population-level evidence about how behavioral state shapes inter-areal coupling in the visual system.

## 2 Population-level GLM coupling model (pop-GLM)

### 2.1 Overview

Point process generalized linear models (GLMs) are widely used for spike train analysis [31; 32; 33]. In these models, the timeline is segmented into sufficiently small time bins to ensure that at most one spike can occur in each bin, and the spike count in each bin is often considered to follow a Poisson distribution. The bin-specific probability of a spike occurring, also known as the instantaneous firing rate, is a latent variable that depends on stimuli, behavior, or past neuronal activity. This firing rate is fitted using generalized regression methods and basis functions, such as splines. By fitting the model, we not only obtain an estimate of the instantaneous firing rate but also can perform statistical inference on various influencing factors. For instance, a single-neuron GLM coupling model focuses on the coupling effects across neurons. In its usual form, it specifies that the log instantaneous firing rate is a summation of three components: (1) stimulus effects, either represented by a time-varying baseline or the convolution of a stimulus filter with the stimulus; (2) self-history effects, commonly represented by the convolution of a post-spike filter with the target neuron’s spike train; and (3) coupling effects, typically represented by the convolution of one or more other source neurons’ spike trains with coupling filters. Through this approach, we obtain an estimate of the functional coupling strength from source neurons to the target neuron while controlling for potential confounding effects such as stimulus and the target neuron’s autoeffects, distinguishing GLM-based methods from pairwise analyses like canonical correlation analysis (CCA) and reduced rank regression (RRR)[5; 34].

We extend this framework to the population level by modeling interactions between pooled population spike trains. In our population-level GLM coupling model (pop-GLM), the spike train for each population is constructed by pooling the spike times of multiple neurons within the same brain area that exhibit similar firing patterns (Figure 1A). Neurons are grouped into populations using a pre-selection procedure based on firing pattern similarity, following the approach of Chen et al. [17]; further details are provided in the Materials and Methods section. For each area, we retain only the population with the strongest stimulus tuning, resulting in a single population per area. As shown in Figure 1B, pop-GLM directly estimates coupling filters between these population spike trains, bypassing the need to fit single-neuron couplings.

**Figure 1:**
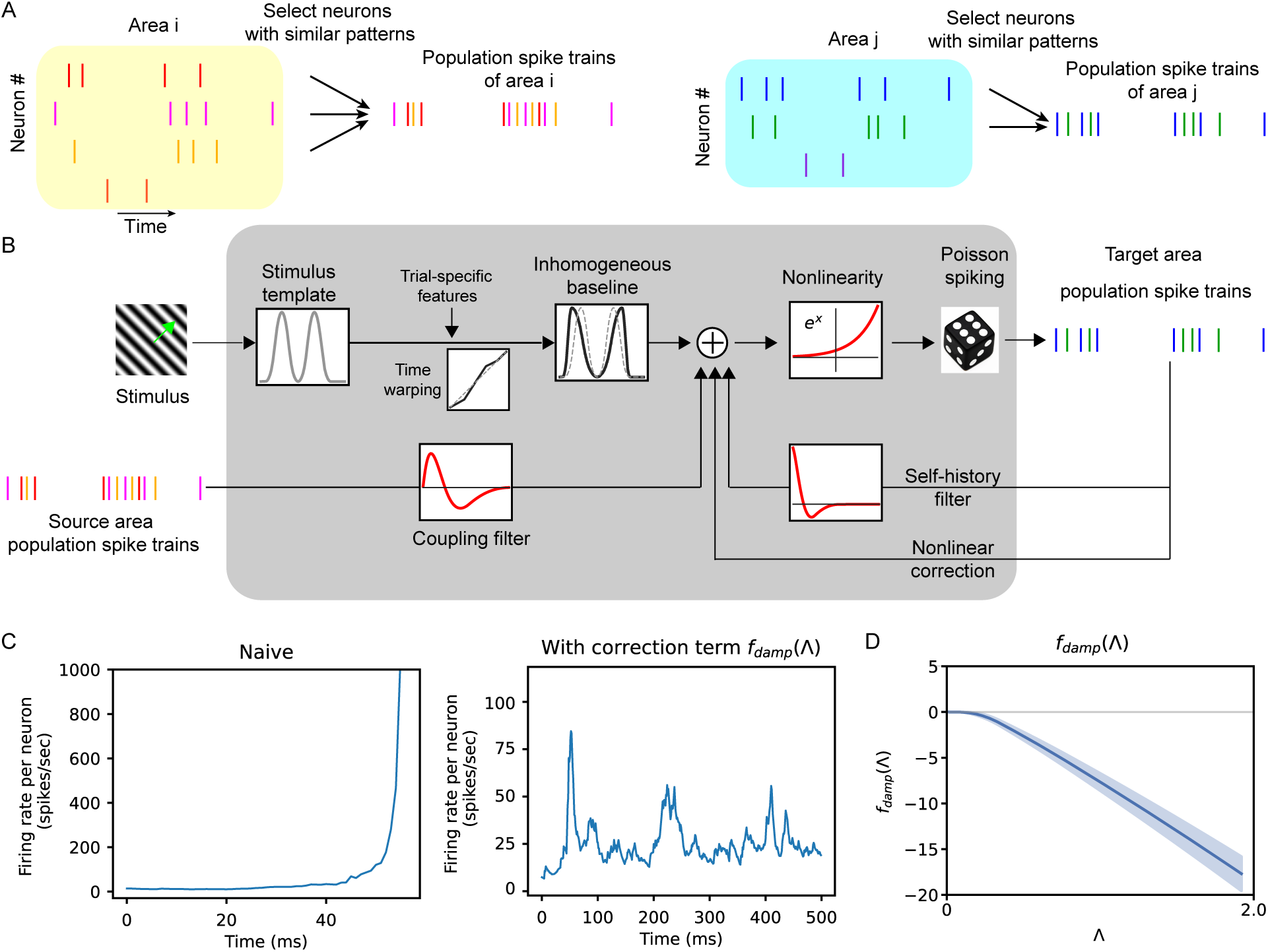
Illustration of pop-GLM. **A**, Population spike trains are obtained by pooling spike trains from a group of individual neurons. **B**, Schematic diagram of the Population GLM Coupling Model. **C**, Comparison of model stability in simulation with and without the correction term *f*_damp_(Λ). After fitting the models with real data, we use the models to do simulations and generate artificial spike trains. On the left, excluding *f*_damp_(Λ) from Equation 1 leads to an unrealistic escalation in simulated firing rate, known as the “explosion” or instability issue. On the right, the introduction of the nonlinear correction term *f*_damp_(Λ) resolves this problem and improves the fit. **D**, Fitted nonlinear correction term *f*_damp_(Λ) of V1 under stationary conditions. This term becomes significantly negative as the empirical firing rate increases. Similar monotonically decreasing trends of *f*_damp_(Λ) are observed in all other areas’ population and conditions, as shown in S12 Fig.

In developing pop-GLM, we introduced two key modifications to the standard GLM formulation to better accommodate population spike trains. First, we explicitly account for trial-to-trial variations in the stimulus effects. This allows us to better account for common drivers and achieve more accurate modeling of population interactions. This trial-to-trial variation is not an automatic consequence of superposition; rather, pooling across neurons enables reliable estimation of trial-to-trial variations that would be difficult to recover from individual neurons alone without further assumptions. Second, we address a stability issue that arises when applying conventional GLM self-history terms to population spike trains. Initial attempts to fit population-level GLMs using standard linear self-history filters resulted in unstable simulations, where predicted firing rates escalated unrealistically (Figure 1C, left). While instability in single-neuron GLMs has been studied and addressed in prior work [35; 36; 37; 38; 39], these solutions do not apply to population spike trains. We found that pooling spike trains suppresses individual refractoriness, making linear history filters insufficient to stabilize the model. To address this, we introduce a nonlinear correction term, *f*_damp_(Λ), which depends on the instantaneous firing rate and acts as a regularizing feedback term (Figure 1C, right and 1D). This correction consistently prevents runaway dynamics and improves simulation fidelity. Detailed descriptions for the two modifications will be provided in Section 2.2 and 2.3.

In schematic form, a standard single-neuron point-process GLM expresses the log firing rate as

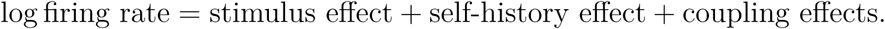

The conditional intensity thus separates into a history-independent component, driven by the stimulus, and history-dependent components, driven by past spiking. The two pop-GLM modifications act on these two parts separately: the history-independent component is made trial-specific, and the history-dependent component acquires a nonlinear damping term. This gives the schematic pop-GLM

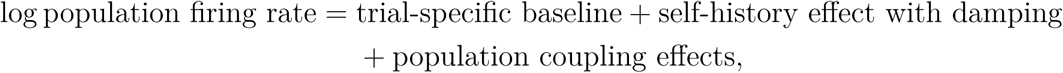

where the “coupling” term captures effects from interactions with other neural populations, and the “self-history” component accounts for local intra-population dynamics. We refer to the history-independent component as the trial-specific baseline throughout.

The trial-specific baseline is further decomposed as:

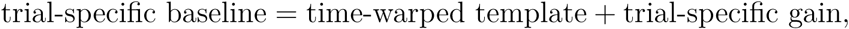

where the time-warped template accounts for shifts in peak response timing across trials, and the gain term captures modulation of overall response amplitude across trials.

The self-history effects are modeled as:

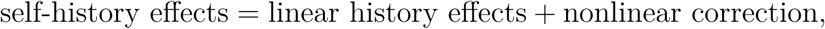

where “linear history effects” is defined as the convolution of the population spike train with a self-history filter, which mainly reflects interneuronal coupling within the population, given that each individual neuron contributes only a minor portion to the total spikes in the population spike trains. The “nonlinear correction” term is more closely related to the refractory effects of each individual neuron.

### 2.2 Modeling trial-to-trial variation in stimulus effects

The stimulus-related term, often referred to as the inhomogeneous firing rate baseline and modeled as a function of time since stimulus onset, is typically assumed to be consistent across repeated trials. However, endogenous states such as arousal or attention, as well as behaviors like locomotion, can modulate this baseline, leading to substantial trial-to-trial variability [40; 41; 42; 43; 44]. In the Allen Brain Observatory – Neuropixels Visual Coding dataset [45], the neural response to a drifting grating stimulus typically exhibits two peaks (Figure 5). Notably, the timing of these peaks, particularly the second one, varies considerably across trials [17; 18].

In fact, the width of the peaks in the PSTH (Figure 5) largely reflects trial-to-trial variability in peak timing. This is evident after applying time warping to shift peak times across trials, which results in a firing rate template that is much narrower than the original PSTH and provides a better fit to the data (S3 Fig). Given these significant variations in neural response, it raises questions about the validity of assuming identical firing rate baseline across all trials. Furthermore, our analysis of a synthetic dataset (Section 3.1) indicates that ignoring trial-to-trial variation in the baseline term can lead to biased estimates of coupling between populations. This bias arises when shared variability in stimulus input acts as a confounding factor; if not properly accounted for, such variability may be misattributed to inter-population coupling. Therefore, it is important to incorporate trial-to-trial variation in the baseline term within the model.

Estimating a unique baseline function for each trial without any constraints is impractical, as it would make the term overly flexible and lead to non-identifiability issues. To effectively capture the most prominent trial-to-trial variability, we instead apply time warping to a baseline function template and introduce a gain constant for each trial. This yields a distinct, yet constrained, baseline function per trial. Time warping accounts for variability in the timing of neural responses, while the gain constant captures differences in overall firing rate, potentially driven by factors such as arousal, engagement, or locomotion [19; 27; 46]. This strategy balances flexibility and identifiability, enabling the model to capture essential trial-to-trial variation without causing non-identifiability.

To capture trial-specific shifts in stimulus responses, we apply a piecewise-linear time warping function *ϕ_j,m_*: [0*, T*] *→* [0*, T*] to the baseline template *f_j_*(*t*), where *T* is the trial duration. This produces a time-warped trial-specific baseline function 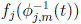 for population *j* on trial *m*. The time warping function is defined using key landmarks that align the two peaks of the average response template to their trial-specific timing (see Section 5.1 for details), allowing the model to flexibly accommodate variability in response timing across trials. Additionally, we introduce a trial-specific gain parameter *β_j,m_* for population *j* on trial *m* to capture differences in overall response magnitude across trials.

### 2.3 Nonlinear self-history effects

In our model, self-history effects are included to capture the influence of recent spiking activity within a population. However, when these effects are modeled solely with a linear convolution term, the fitted model becomes unstable when using them to generate spike trains, resulting in a rapid and unrealistic escalation in firing rate (Figure 1C). This phenomenon, commonly known as the “explosion” problem in GLMs, has been extensively studied in single-neuron settings, where various causes and remedies have been proposed [35]. However, in the population-level scenario addressed here, the solutions applicable to single-neuron GLMs fail to resolve this problem.

We found that the pooling procedure is sufficient to cause the explosion issue. From a neurophysiological standpoint, the refractory period sets a limitation for firing rate of neurons. When modeling the firing rate of single neurons with GLMs, we often observe a pronounced negative dip at the beginning of the self-history filter. This dip reflects the strong refractory effects. Consequently, the firing rate drops to almost zero immediately after a previous spike. However, when modeling the population firing rate, with self-history effects assumed to be linear, the extent of refractory effects is greatly underestimated or even disappears. We support this point by both analytical and empirical results in S1 Text. Consequently, the system lacks a “brake” to counteract positive feedback loops, which can drive the system to a firing rate that is unrealistic, as illustrated in S1 Fig.

To approximate refractory effects that are not captured by the linear self-history term, we introduce a nonlinear correction term *f*_damp_(Λ), where 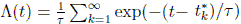 represents a smoothed count of recent spikes within an exponential window. Here, 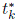 denotes the time of the *k*-th most recent spike before time *t*, and the hyperparameter *τ* controls the width of the temporal window. The function *f*_damp_ is represented as a weighted sum of basis functions of the recent population spike density Λ(*t*); specifically, we parameterize it with four raised-cosine basis functions spaced over the range of Λ (S4 Fig, panel C). The fitted function is consistently negative, as shown in Figure 1D. This provides dynamic negative feedback when excessive spiking occurs in a short time interval. The correction term not only stabilizes the model by preventing runaway firing rates, but also improves the fit. We tested alternative forms of nonlinearity in the self-history component, but found that *f*_damp_(Λ) yielded the best overall performance (Table A in S1 Text).

One potential concern is that the nonlinear correction term may not fully capture the true history effects, potentially introducing bias into the estimated coupling filters. To assess this, we re-estimated the coupling filters using exact self-history terms from individual neurons, replacing the correction term *f*_damp_(Λ). As shown in S7 Fig, the resulting coupling filters remain largely unchanged; quantitatively, across all 60 cross-area coupling filters the two models agree closely in both amplitude (correlation of filter integrals *r* = 0.93) and timing (correlation of peak latencies *r* = 0.81), and the V1–LM result is preserved. This underscores the efficacy and relative simplicity of the correction term as an effective approximation.

### 2.4 Pop-GLM equations

We formally define our population-level GLM (pop-GLM) in Equation 1, which models the log firing rate of a population spike train:

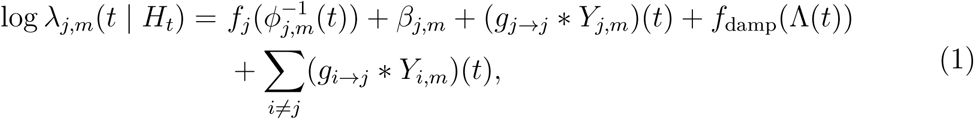

where *λ_j,m_*(*t | H_t_*) denotes the instantaneous firing rate of the population spike train for population *j* at time *t* on trial *m*, conditioned on the spike history *H_t_*. The term *f_j_*(*t*) is the inhomogeneous baseline template for population *j*. The trial-specific time warping function *ϕ_j,m_*(*t*) shifts the timing of the two peaks in *f_j_*(*t*) for trial *m*, and the gain constant *β_j,m_* scales the overall response magnitude for trial *m*. Together, 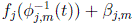 defines the trial-specific inhomogeneous baseline for population *j*.

The term *Y_j,m_* denotes the population spike train in area *j* on trial *m*. The self-history filter *g_j_*_→_*_j_* captures internal temporal dependencies within population *j*, while *g_i_*_→_*_j_* represents coupling from source population *i* to target population *j*. The nonlinear correction term *f*_damp_(Λ(*t*)) modifies the self-history effects, where 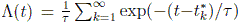 captures recent spike activity through an exponentially weighted sum over past spike times 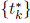. This term helps to account for refractory effects and prevents unrealistic escalation of firing rates during simulations.

Each component of the model is parameterized using basis functions to enable flexible and interpretable estimation. The baseline template *f_j_*(*t*) is modeled using a B-spline basis. The coupling filters *g_i_*_→_*_j_*(*t*), the self-history filter *g_j_*_→_*_j_*(*t*), and the nonlinear correction term *f*_damp_ are parameterized using a set of raised cosine basis functions as in Pillow et al. [33]. All basis coefficients are estimated via maximum likelihood. The time warping functions are fitted greedily in alternation with these regression coefficients as part of the optimization procedure.

The central focus of our model, the cross-population coupling filter *g_i_*_→_*_j_*(*t*), remains interpretable as in single-neuron GLMs: it represents the change in the log firing rate of the population spike train for population *j* following a spike in population *i* with a time lag of *t*. Since the log firing rate of a population spike train differs from that of the mean single-neuron firing rate by a constant equal to the logarithm of the number of neurons, this filter can also be interpreted as the effect on the mean firing rate of individual neurons in population *j*.

## 3 Results

### 3.1 Results with synthetic dataset

#### Simulation setup

Because the ground truth is unavailable in real data, we began by evaluating our population-level GLM coupling model (pop-GLM) on synthetic datasets, comparing it against existing single-neuron GLMs [9; 10] and reduced-rank regression (RRR) [34]. While RRR does not capture conditional dependencies like the GLM-based methods, we include it for completeness. We simulated spike trains using exponential integrate-and-fire (EIF) neurons [13; 47; 48] to construct a source and a target neural population, designed to mimic the V1 and LM activity observed in our real data. The model includes mutual excitation within each population using synaptic weights drawn from a log-normal distribution [49]. Simulation details are provided in the Materials and Methods.

#### Scenario 1: Detecting weak coupling

We consider two scenarios. The first tests the sensitivity of each method in detecting weak coupling from the source to the target population. Both populations receive trial-invariant input, with weak synaptic connections from source to target drawn from a log-normal distribution (Figure 2A). Figures 3A show the estimated coupling filters when we fit pop-GLM and single-neuron GLMs to the simulated spike trains. It is evident that while the coupling filter estimated by single-neuron GLMs shares the same expectation, it exhibits significantly higher variance than those fitted by pop-GLM. This discrepancy arises because single-neuron GLMs involve a greater number of parameters (scaling quadratically with the number of neurons in a population), thereby increasing model variance. We then conducted a likelihood ratio test on the three models to test the null hypothesis of no coupling between the two populations. With 100 trials, approximately the number in the real dataset, pop-GLM demonstrates greater sensitivity in detecting coupling, shown by its significantly higher area under the curve (AUC) (Figure 3B). For other trial counts, as the number of trials increases, all three methods yielded smaller *p*-values against the null hypothesis (Figure 3D) and exhibit higher statistical power (Figure 3C). Among the three methods, pop-GLM is the most sensitive, requiring only about 100 trials to achieve a statistical power of 0.8 (rejecting the null hypothesis 80% of the time) at a significance level of 0.05, whereas the single-neuron GLM needs approximately 250 trials. Reduced rank regression, which neither takes advantage of Poisson likelihood nor can use smaller time bins (requiring a 50 ms bin and thus limiting its effective sample size), performed worst and requires more than 800 trials to reach a power of 0.8.

**Figure 2:**
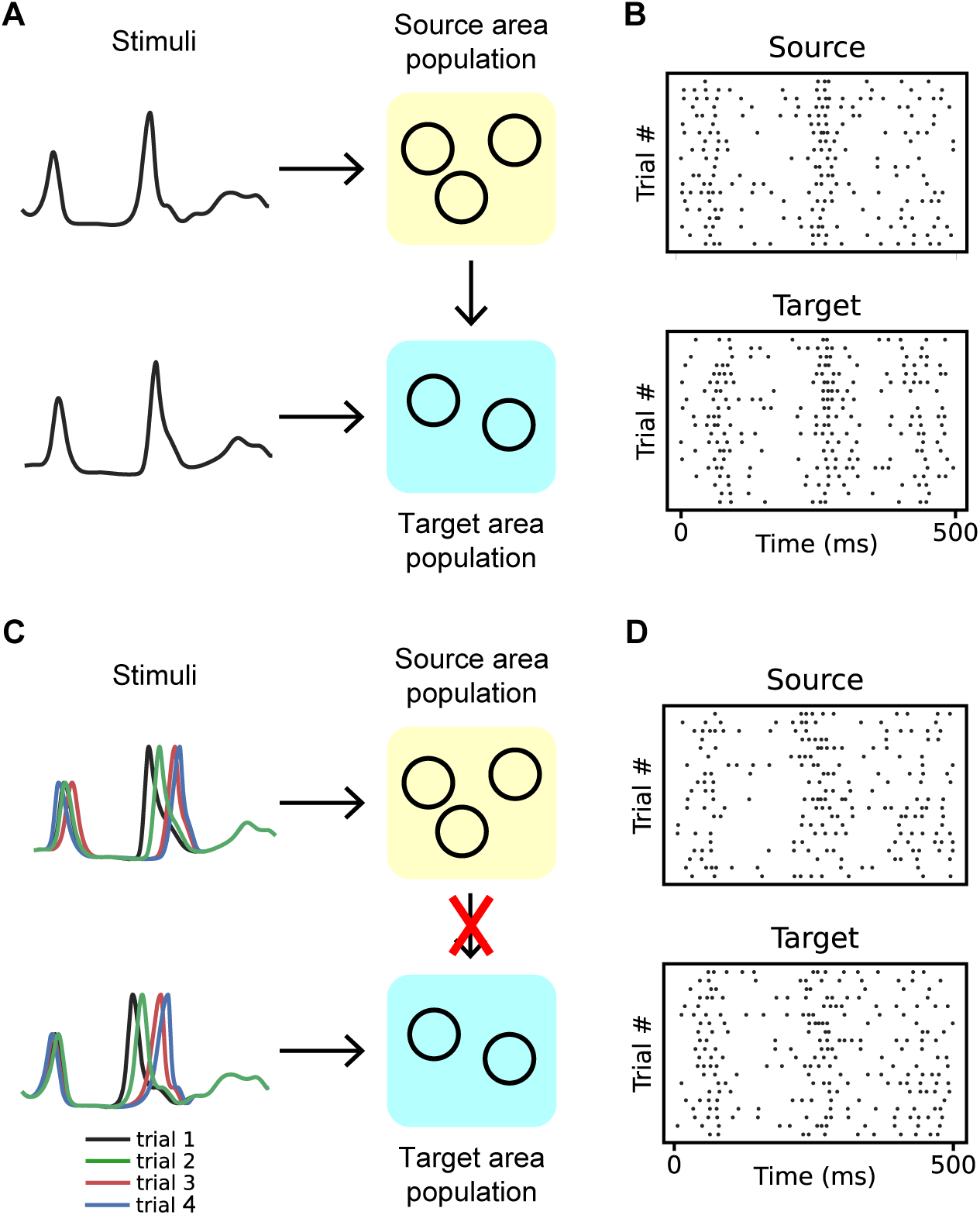
Illustration of two scenarios in synthetic datasets. **A&B**, The first scenario. **A**, We created two populations, a source population and a target population, each comprising ten exponential integrate-and-fire (EIF) neurons. The external inputs to both populations, which are the same for each trial, are depicted in panel A. Each neuron within a population receives identical input, designed to mimic the firing rates of V1 and LM under stationary conditions (see Figure 5). Additionally, there are very weak synaptic inputs from each neuron in the source population to each neuron in the target population, with the same synaptic weights for each neuron pair. **B**, Spike trains of an example neuron in the source and target population in the first scenario. **C&D**, The second scenario. **C**, The same two populations as in A, but with NO coupling between them. Both populations receive inputs with trial-to-trial variations, featuring two peaks that shift across trials. **D**, Spike trains of an example neuron from both the source and target populations in the second scenario.

**Figure 3:**
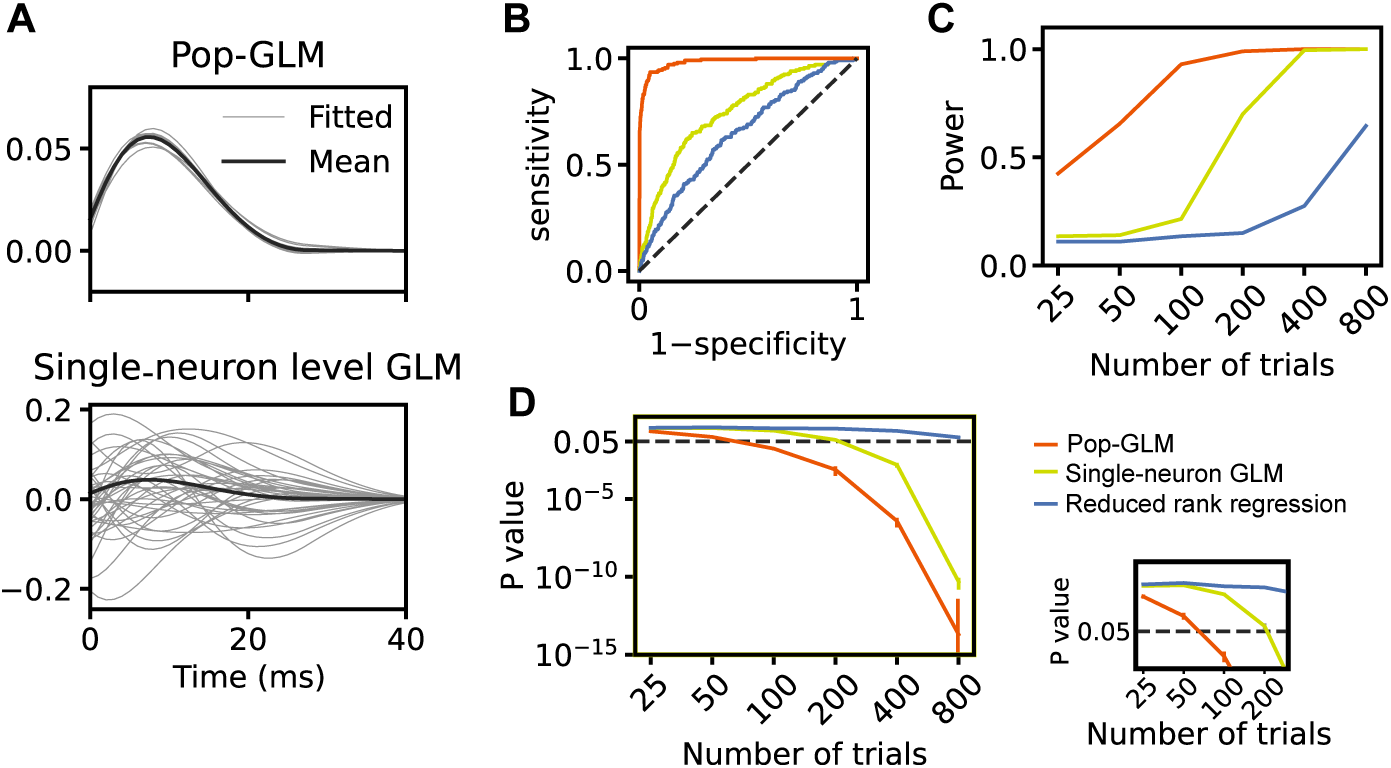
The population-level model requires a much smaller sample size compared to the single-neuron-level model to achieve the same level of sensitivity in detecting cross-population interactions. **A**, Coupling filter fitted by pop-GLM (upper) and single-neuron-level GLMs (lower). The gray curves show the fitted coupling filters across ten repetitions. Filters fitted by single-neuron-level GLMs exhibit significantly higher variability. **B**, The Receiver Operating Characteristic (ROC) curve demonstrates the superior ability of pop-GLM (red) in detecting coupling effects from the source to the target population, compared to single-neuron GLMs (yellow) and reduced rank regession (blue). For pop-GLM, the likelihood ratio test is conducted by fitting a nested model, where the coupling filter from the source to the target population is assumed to be zero, and a full model with no constraints. For the single-neuron-level model, the same likelihood ratio tests are performed for each neuron, and Fisher’s method is used to aggregate these *p*-values into a final *p*-value. For reduced rank regression, a permutation test is conducted by shuffling the source population spike trains across trials, using mean squared error as the test statistic. **C**, The averaged *p*-values (blue) and statistical power (red) for pop-GLM (solid lines) and single-neuron-level GLM (dashed lines) were compared across varying numbers of trials (25, 50, 100, 200, 400, 800), with statistical power measured at a significance level of 0.05. For each trial setting, two hundred repetitions were conducted. Pop-GLM consistently reports lower *p*-values and achieves higher statistical power.

We also explored a wide range of hyperparameters, including the number of neurons in a population, mean synaptic weights, and variance of synaptic weights, and found that the results remain consistent. Notably, when the number of neurons in each population increases, or when we increased the variance of synaptic weights, pop-GLM demonstrated an even greater advantage over single-neuron GLMs. This occurs because increasing the number of neurons leads to a quadratic increase in the number of parameters in single-neuron GLMs, making the fitted coupling filters noisier and thereby reducing statistical power. Similarly, higher variance in synaptic weights also leads to noisier fitted coupling filters, diminishing the statistical power of single-neuron GLMs.

#### Scenario 2: Avoiding false positives under correlated inputs

In the second scenario, there is no ground truth coupling between the populations. However, the input stimulus exhibits trial-to-trial temporal shifts, creating the potential for spurious coupling due to shared variability (Figure 2B). We configured the stimulus input to mimic the activity in V1 and LM as observed in the real dataset, where the two peaks are temporally shifted on a trial-by-trial basis (Figure 3A). The correlation between the peak times of the source and target populations also aligns with the real data correction ([17] Figure 1 B and C). This highly correlated stimulus input can act as a confounder in the relationship between the two populations. This was observed in Figure 4B and C, where, without accounting for the trial-to-trial variation in stimulus input, both the population-level model and the single-neuron model yield very small *p*-values, leading to the false rejection of the null hypothesis. However, the population spike trains contain sufficient information to estimate these shifts using time warping. As a result, our pop-GLM with time-warping equipped consistently does not reject the null hypothesis. In contrast, single-neuron GLMs lack enough information per neuron to estimate per-trial variability, making some form of neuron pooling effectively necessary. This highlights that modeling trial-level stimulus variability is inherently a population-level problem.

**Figure 4:**
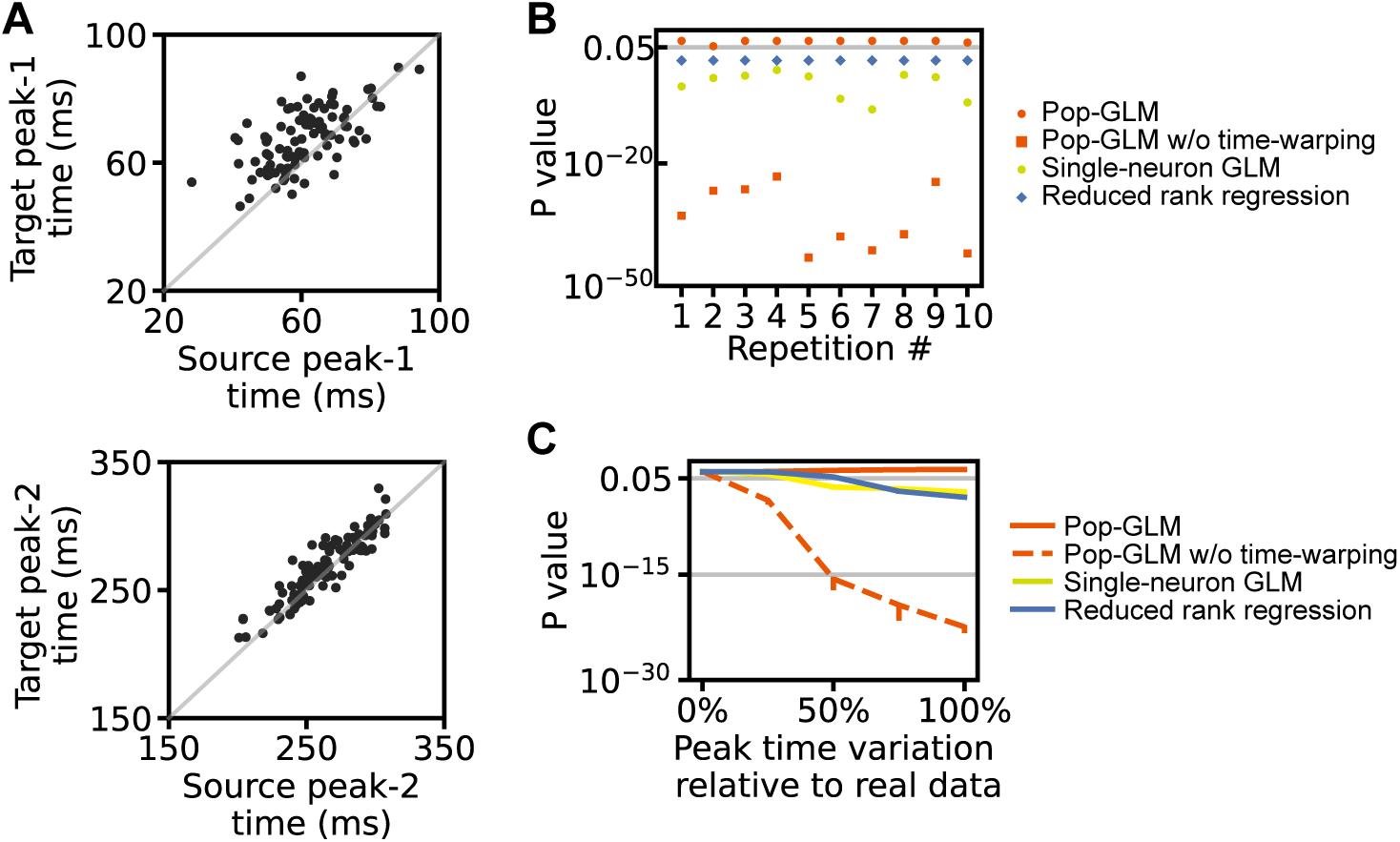
The population-level model with a time-warping component to account for trial-to-trial variation effectively avoids false discoveries in cross-population interactions influenced by highly-correlated backgrounds. **A**, The timing of peak-1 (upper) and peak-2 (lower) in both populations. Peak-1 times show a correlation of 0.5, and peak-2 times, a correlation of 0.9, consistent with V1 and LM results from real data. Their standard deviations and means also match those found in real data. **B**, Inference was conducted similarly to Figure 3, but in this case, there is no ground truth connection between the two populations. Without time-warping component, single-neuron models (yellow dots), reduced rank regression (blue diamonds), and population-level models (red squares) would mistakenly detect coupling between the populations with very small *p*-values. Pop-GLM equiped with time-warping (red dots) remains robust against false positives in this scenario. **C**, An increase in the standard deviation of peak times causes models without time-warping (single-neuron models: gray solid; population-level models: black dashed) to report increasingly significant false positive results. In contrast, pop-GLM (black solid) continues to demonstrate robustness. On the x-axis, a 0% value indicates no variation, while 100% reflects the variation matching that of the real data.

**Figure 5:**
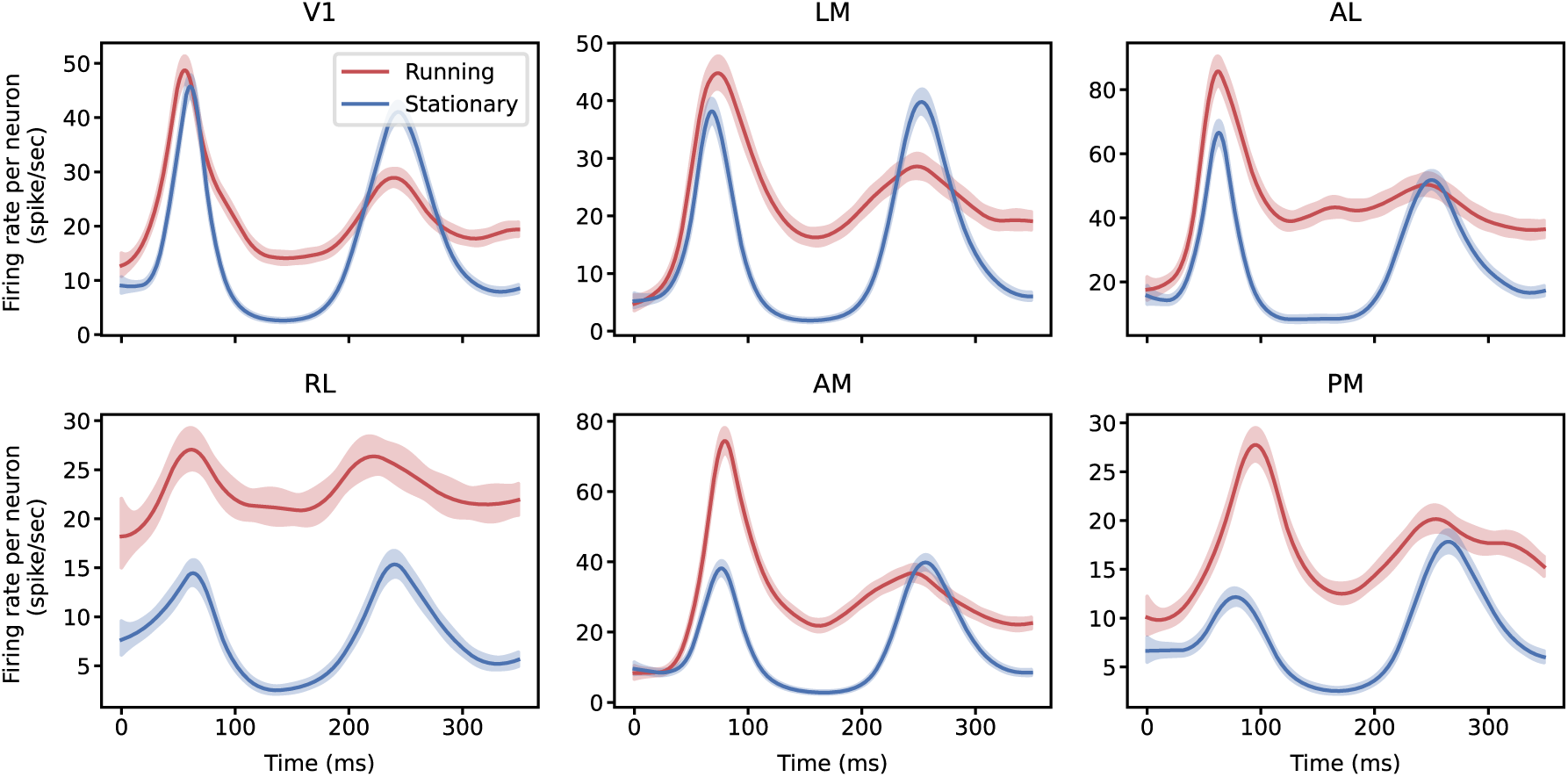
Neural population response to the first 350 ms of a full-field drifting grating stimulus. The stimulus begins at 0ms. The mean firing rates of six populations in six visual areas, under running and stationary conditions, are fitted using smoothing splines with second derivative penalties. The first peak occurs around 60 ms, followed by a second peak around 250 ms. When the mouse is running, the overall firing rate is higher, whereas the second peak is more prominent in the stationary condition. The drifting grating stimulus begins at 0 ms. There are a total of 129 and 81 trials for stationary and running conditions, respectively, across 14 stimulus conditions.

#### Scenario 3: Coupling and trial-to-trial variability together

Scenarios 1 and 2 isolate the two effects: coupling with no trial-to-trial variability (where the fitted filter should be nonzero), and trial-to-trial variability with no coupling (where it should be zero). The realistic situation is that both are present at once, and one might worry that the time-warped baseline could absorb genuine coupling-related structure. We therefore simulated the same EIF populations with *both* a nonzero feed-forward coupling and across-trial jitter in the timing of the two response peaks, keeping all other settings as in Scenario 1, and fitted pop-GLM with and without the time-warping component, along with the single-neuron GLMs (S13 Fig).

Three results follow. First, pop-GLM with time warping recovers a *nonzero* coupling filter whose size matches the filter recovered in the absence of any trial-to-trial variability (integral 0.74 *±* 0.12 versus a no-variability reference of 0.75 *±* 0.17). The time-warped baseline therefore preserves true coupling rather than absorbing it. Second, removing the time-warping component from pop-GLM inflates the fitted filter: the unmodelled trial-to-trial variation is misattributed to coupling. Third, the filters fitted by single-neuron GLMs are not only far noisier than pop-GLM’s but are also inflated to the same degree as the no-time-warping pop-GLM estimate. These results further support the robustness of our conclusions from Scenario 1 and Scenario 2.

#### Summary

The synthetic data results highlight two key advantages of pop-GLM. First, when weak coupling is present, pop-GLM detects connectivity with greater sensitivity and requires fewer trials than competing methods. Second, when correlated trial-to-trial variability exists in the inputs to both populations, modeling baseline shifts through time-warped templates, made possible by pooling across neurons, helps avoid false positives. Together, these results demonstrate that population-level modeling enhances statistical power and reduces bias caused by ignoring trial-to-trial variability in input structure.

### 3.2 Results with experimental data

#### Stimulus response in six visual areas

Before presenting the fitted results of pop-GLM, we first show the average evoked responses to full-field drifting grating stimuli across six visual areas (V1, LM, AL, RL, AM, and PM). The stimulus is presented at time 0, and neural responses typically exhibit two peaks around 60 ms and 250 ms. This dual-peaked sensory response has been previously reported in early visual areas [50; 51; 52]. The first peak is thought to reflect feedforward propagation of sensory input, while the second peak is modulated by top-down feedback and has been associated with perceptual processing [17; 51; 52]. We designed our synthetic dataset to closely mimic this dual-peaked neural activity in the real dataset, and our hand-designed trial-to-trial variation with time-warping is also designed to capture the most important aspect of the dual-peaked response.

Although two peaks are visible in all six areas under both conditions, the firing patterns are markedly different between running and stationary conditions (Figure 5). The overall firing rate is higher when the mouse is running, consistent with previous findings [19; 20; 21; 23; 24; 25; 26; 27]. Furthermore, the second peak in the running condition is notably less prominent than that in the stationary conditions. This observation leads us to the central question of our study: does cross-area coupling also change with these behaviorally modulated firing dynamics?

#### Fitted coupling filters

We fitted pop-GLM to both running and stationary trials separately and used permutation tests to find differences in coupling filters between the two conditions. A trial was classified as running or stationary by thresholding its average running speed at 1 cm/s, a rule selected by comparing alternative ways of incorporating speed into the model (Table B in S1 Text). Figure 6 shows all coupling filters from one area to another, along with the corresponding baseline firing rate templates. After correcting for multiple comparisons, we found that the excitatory coupling filter from V1 to LM was significantly attenuated during locomotion (*p* = 0.0014 after Bonferroni correction; *p* = 0.0011 with a more precise permutation-based correction procedure described in the Statistical testing section). A spike in V1 is associated with 0.05 additional spikes in LM, conditioning on all other terms in the model, when the mouse is stationary, but this drops to 0.04 during running. Coupling from RL to AM and from V1 to other higher-order areas such as AL and RL also appears reduced during locomotion, though their *p*-values did not reach significance after correction.

**Figure 6:**
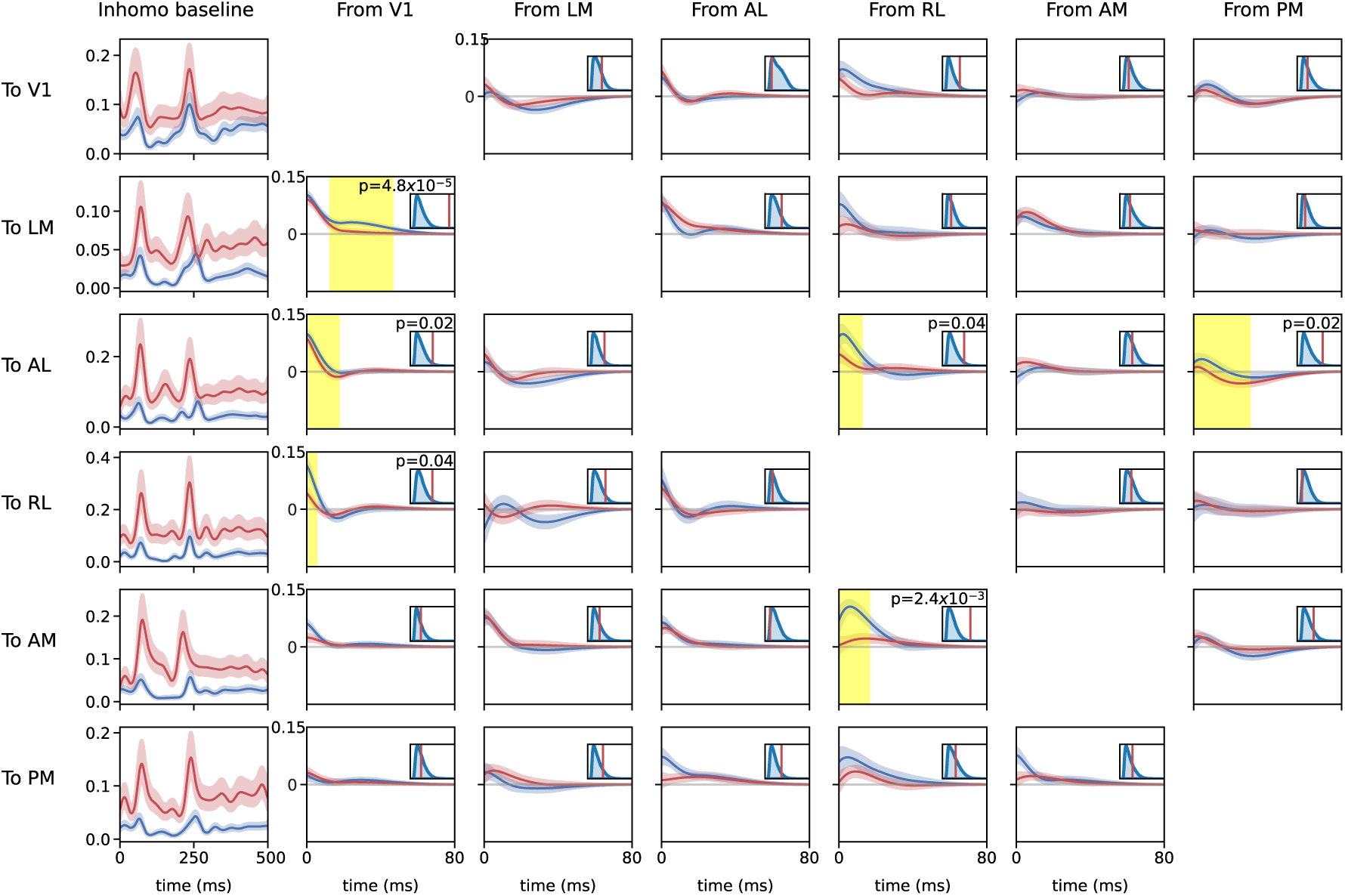
Fitted GLM coupling filters and inhomogeneous baseline firing rate templates (95% CI). Red: running. Blue: resting. The smaller figure in the upper right of each panel displays the results of excursion tests, where the blue distribution represents the excursion test statistic distribution under the null hypothesis, and the red line indicates the observed statistic. The raw *p*-values are also shown in the figure. Yellow areas highlight regions of interest from excursion tests with *p*-values below 0.05, prior to correcting for multiple comparisons. The six panels of each row depict different components of a GLM model. For example, the panels in the first row, from left to right, present the inhomogeneous baseline for V1, the coupling filter from LM to V1, the coupling filter from AL to V1, the coupling filter from RL to V1, the coupling filter from AM to V1, and the coupling filter from PM to V1, respectively. Since we have a gain constant for each trial, the effects of different numbers of neurons in each area are mitigated, and coupling filters are normalized to spikes per second per neuron scale.

To further validate our findings, we replicated the model fitting and inference process with a second mouse, without changing any hyperparameters. Again, we observed significant attenuation of V1-to-LM coupling during running (Figure S11). Since we tested only this connection in the second mouse, no multiple comparisons correction was required.

#### Similar findings using LFP data

Local field potential (LFP) and spiking data, both signals from extracellular recordings, differ substantially in their physiological origins. While spiking data specifically reflects the firing activity of individual neurons close to the electrode, LFPs primarily capture the collective synaptic activities of a large number of neurons in proximity to an electrode. Analyzing the LFP data also revealed two notable peaks, closely matching the timing of two firing rate peaks in the spike trains (Figure S10A). We applied Granger causality to assess functional connectivity in LFP data, and observed a reduced connection from V1 to LM during locomotion (Figure S10B). We replicated this LFP analysis with the second mouse, and the results were consistent.

#### Failure of single-neuron GLMs

To directly compare against pop-GLM, we applied single-neuron GLM to the same dataset. The fitted coupling filters and coupling outputs are shown in Figures S8 and S9. However, the single-neuron GLMs yielded implausible results, most notably a pervasive inhibitory influence from V1 onto all other areas. The inhibition sent from V1 was markedly strong, except when V1 was in the period preceding stimulus onset and in the firing rate dip between the two peaks. These two periods of reduced activity in V1 actually correspond to the two firing rate peaks in other areas, considering an approximate 40 ms delay in the inhibition effects. In this way, the model essentially learns a baseline function with trial-to-trial variability and fits the data better. However, this pattern of consistently strong inhibition is more likely an artifact, potentially stemming from the model’s inability to account for trial-to-trial variability in the stimulus effect. These results underscore the limitations of single-neuron GLMs and highlight the importance of modeling trial-level population dynamics, as done in pop-GLM.

#### Suggested dynamics of the fitted model

Connections between generalized linear models (GLMs) and mechanistic models, such as the leaky integrate-and-fire (LIF) model, are established [53; 13]. In particular, the log firing rate in a GLM can be qualitatively interpreted as a proxy for membrane potential in an LIF model, and the fitted components of GLMs can reflect underlying currents or voltage changes.

Viewed through this lens, our fitted GLM exhibits dynamics consistent with a simple dynamical model. Figure 7 shows the average contribution of each component in our fitted model on LM; the trial-averaged fitted outputs of every component across all six areas are shown in S14 Fig. The self-history effect of LM itself acts as a strong driver, in addition to the input from V1 after the initial stage of sensory processing. This result aligns with the assumption of strong recurrent excitation [54; 55; 16].

**Figure 7:**
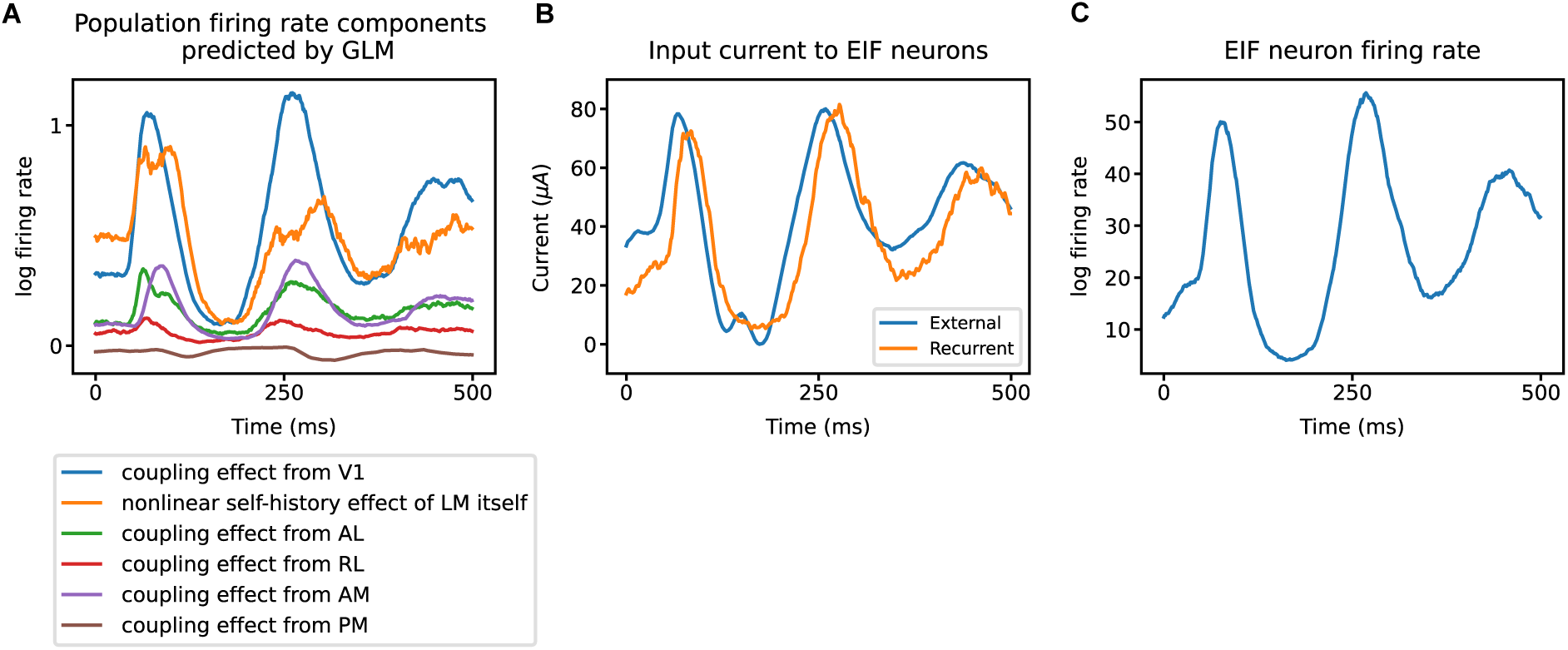
Strong recurrent excitation suggested by pop-GLM. **A**, Each component of LM intensity, averaged across trials, predicted by pop-GLM. The blue curve represents the input from V1. The orange curve represents the nonlinear self-history effect of LM itself, consisting of both the term *f*_damp_(Λ) and the output of the linear post-spike filter. The other four curves are the linear coupling effects from the rest four areas. The inputs from V1 and from LM itself are the two biggest drivers. **B**, Trial-averaged current to ten EIF neurons mimicking the LM population. For each trial, the external input (blue) is the sum of all inputs predicted by the GLM, except those from LM itself. There are also recurrent connections within the ten-neuron population, and the orange line shows the trial-averaged input from other neurons within the population. **C**, The EIF neuron population generated spike trains with firing rates similar to those observed experimentally.

To further examine this interpretation, we simulated a recurrent population of ten EIF neurons driven by external input. The external input to the EIF neuron population was a rescaled sum of all fitted components of pop-GLM, except for the nonlinear self-history effect, which was replaced by recurrent connections between these neurons. This configuration of the EIF neuron population was able to generate a PSTH similar to those observed experimentally (S5 Fig), with the self-excitation within the population strongly following external input (shown in panels B and C of Figure 7). These findings support the view that the fitted pop-GLM effectively captures dynamics consistent with those of a recurrently connected EIF population.

## 4 Discussion

Because recent work obtained interesting results by focusing on small populations of neurons that exhibited homogeneous responses to a specific stimulus [17; 18], we adopted a population firing rate perspective in the spirit of Chen et al. [17], and developed a population point process model. We explored the utility of point process GLM coupling models at the population level and, using synthetic data from recur-rent EIF models, we found that these population models can be more sensitive in detecting coupling across areas than aggregating effects from individual neuron models. Population-level models also allowed us to consider more easily the trial-to-trial variation in response to a stimulus, which appears to be a major reason pop-GLM produced more intuitive results than an aggregated individual-neuron procedure when applied to the visual cortex spiking data we analyzed.

Using our pop-GLM method on experimental data, we observed in two mice that functional connectivity from V1 to LM is weakened during locomotion. Our analysis of LFPs also supported this finding. Previous work has demonstrated that locomotion drives an overall increase in firing rates in visual areas, a finding which we replicate here. In light of the present work, it appears this increased activity level does not co-occur with increased coupling between V1 and higher visual areas. This suggests that firing rate increases do not result from more effective propagation of spikes from the periphery (mediated by V1), but rather from a modulation of overall activity levels by a shared input. Whether this originates from motor feedback (e.g., efference copy), neuromodulatory signals, or some other source is currently unclear.

We also wish to address several limitations of our work. (1) Although we used a second mouse to check the consistency of our results, examining additional subjects and quantifying the degree of subject-to-subject variation in V1 to LM functional connectivity could yield more definitive conclusions. (2) Additionally, exploring how coupling changes in response to different types of visual stimuli, as well as to variables correlated with global states such as arousal, would provide further insights. (3) Nonlinear spike-history effects effectively prevent instability and improve the fit; however, our nonlinear correction term was somewhat ad hoc, challenging to interpret, and could potentially be improved. (4) Lastly, we assumed that population coupling effects re-main constant within the time interval analyzed for each trial, though the temporal evolution of functional connectivity deserves further investigation.

Separately from these specific points, it is worth stating explicitly how pop-GLM would be expected to behave in regimes beyond the strongly stimulus-locked cortical populations studied here. Two parts of the model are separable: the pooled coupling formulation, which is the source of the gain in statistical power, and the trial-specific baseline, which is what requires a repeated, landmark-identifiable stimulus response. For heterogeneous populations, our grouping procedure assumes that neurons within a population share a common functional response; the pooled spike train remains well defined when they do not, but the interpretation of the fitted filter changes, and it should then be understood as an average coupling effect across the pooled neurons rather than as a property shared by each of them. If a group mixes neurons whose cross-area interactions differ in sign, excitatory and inhibitory contributions can partially cancel and the population filter will understate both. In this case, it would be best either to divide an area into finer sub-populations before fitting, or to interpret the filter explicitly as a population average. Spontaneous activity, weakly evoked responses, and tasks without a repeated stimulus template challenge the second part of our model: the landmark-based warping of the trial-specific baseline has nothing to align to. The rest of the model remains well posed and the gain in statistical power is untouched; the risk is that slow shared fluctuations driven by arousal or movement, no longer locked to trial structure, are misattributed to coupling at long lags. The template should therefore be replaced rather than simply omitted, and because it enters the log firing rate as an additive term it can be replaced without altering the coupling formulation: a piecewise-linear or Gaussian Process trial-specific baseline, or an even more flexible baseline learned from the data [56] rather than specified by hand would serve this role. We therefore regard these regimes as calling for the substitution of a component rather than as obstacles to the approach.

We would like to highlight a final point, one which has been discussed extensively, especially under the general headings of replication and “p-hacking” [57**?**], yet often gets little attention within individual articles. In our reported results here, correction for multiple comparisons was straightforward (though we used a permutation procedure to check the Bonferroni correction, as it can be conservative). More worrisome are the many preliminary steps that precede formal analysis because they involve pre-processing and feature definition, which are not accounted for when computing assessments such as *p*-values. In analyzing locomotion, an obvious consideration is the definition of “running”. If the way we defined running somehow were to depend on the coupling effects we ended up examining (for example, in an extreme case, if we were to choose a definition that provided improved results), then, obviously, the computed *p*-value would lose its meaning. The models considered in choosing our definition, however, did not include the key variables of interest, namely the coupling terms. Furthermore, to guard against other, unknown potential sources of inferential corruption we replicated the results on a second mouse, repeating neither the relevant pre-processing steps nor the fitting of hyperparameter values. Although when replicating a finding in this way it remains possible that some source of bias was also replicated, the other common concerns are greatly mitigated.

## 5 Materials and methods

### 5.1 Time warping function

To account for trial-to-trial variability in response timing, we apply a piecewise-linear time warping function *ϕ_j,m_*(*t*): [0*, T*] *→* [0*, T*] to the stimulus template *f_j_*(*t*), where *T* = 500 ms is the trial duration. The warped input function is then given by 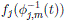, which shifts input peaks on a per-trial basis.

The warping function *ϕ_j,m_* is defined by a set of fixed and data-driven landmark pairs: 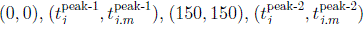, and (500, 500). Here, 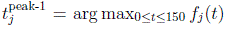 and 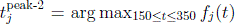 are the first and second peak locations in the template *f_j_*(*t*), while 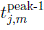 and 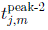 are their trial-specific location after shifting.

This procedure shifts the peaks of the template *f_j_*(*t*), resulting in the time-warped input function 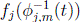. To further model variability in response magnitude across trials, we include a trial-specific gain term *β_j,m_* for population *j* in trial *m*.

### 5.2 Fitting and training procedure

We estimate the pop-GLM parameters by alternating between two steps: (i) penalized maximum-likelihood estimation of all basis coefficients given the current trial-specific time warping and gains, and (ii) greedy updates of the trial-specific warping landmarks and gains given the current coefficients. The full procedure is summarized in Algorithm 1.

The baseline template *f_j_*(*t*) is initialized from the trial-averaged peristimulus time histogram (PSTH), fitted with the B-spline basis; the warping functions are initialized to the identity (landmarks at the template peak times) and the trial gains to *β_j,m_* = 0. In the regression step, the warped baseline, coupling, self-history, and *f*_damp_ bases are assembled into a single design matrix, and all basis coefficients together with the per-trial gains *β_j,m_* are estimated jointly by penalized maximum likelihood using Newton–Raphson (iteratively reweighted least squares) with an *L*_2_ ridge penalty; the inhomogeneous-baseline coefficients are left unpenalized. In the warping step, for each trial *m* the peak-1 and peak-2 landmarks of the warping function *ϕ_j,m_* are updated by a line search that maximizes that trial’s Poisson log-likelihood given the current coefficients, and the landmark updates are damped across iterations with an exponential moving average for stability. The nonlinear damping term *f*_damp_ is updated together with the other coefficients in the regression step. The two steps alternate until the relative decrease in the total negative log-likelihood falls below a tolerance *ɛ* or a maximum number of iterations *M* is reached.

### 5.3 Synthetic dataset simulation

We used exponential integration-and-fire (EIF) neurons [48; 13; 47] to generate the synthetic dataset. Unlike the simpler leaky integrate-and-fire (LIF) model, the EIF model accurately represents the exponential increase in membrane potential near the firing threshold and is considered to have superior capability in mimicking real neuronal dynamics [48]. The spiking neural network we created in our synthetic dataset consists of two neural populations, the source and the target, each comprising ten neurons. The dynamics of the neuron membrane potential is described by the following equation:

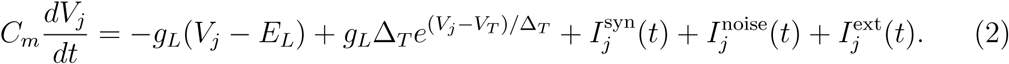

When the membrane potential *V_j_*(*t*) reaches a threshold *V*_th_, the neuron produces a spike. Subsequently, the membrane potential is held for a refractory period *τ*_ref_ and then resets to a fixed value *V*_re_. We use the same parameters as those in Huang et al. [47]: *τ_m_* = *C_m_/g_L_* = 15ms, *E_L_* = *−*60mV, *V_T_* = *−*50mV, *V*_th_ = *−*10mV, Δ*_T_* = 2mV, *V_re_* = *−*65mV, and *τ*_ref_ = 1ms. 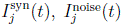 and 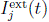 represent synaptic input from other neurons, random input, and the external stimulus.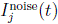 is assumed to be Gaussian white noise. We selected a noise amplitude that makes neurons fire at a 10Hz baseline in the absence of any external stimuli or synaptic inputs. 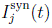 is described by

**Algorithm 1:**
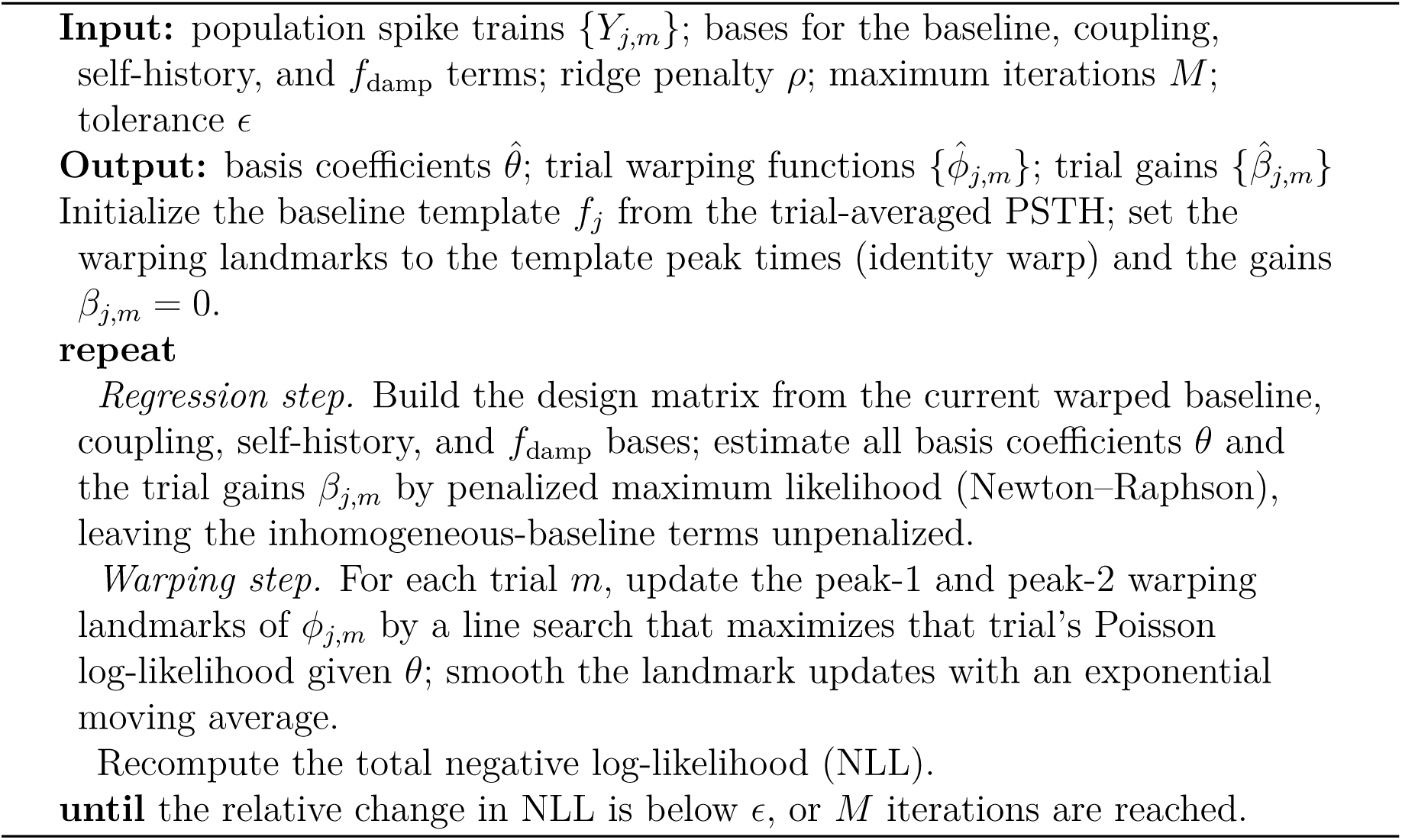
Fitting pop-GLM.

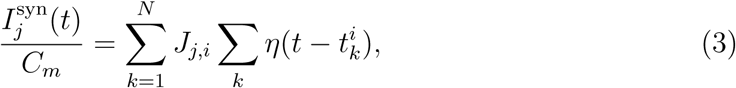

where *J_j,i_* is the synaptic weight from the i-th neuron to the j-th neuron, and 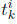 represents the time of the i-th neuron’s k-th past spike. *η* is the unit postsynaptic current, which is given by

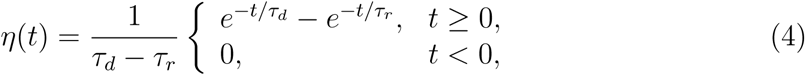

where we use *τ_d_*= 5ms and *τ_r_*= 1ms.

We designed two scenarios to evaluate the performance of our pop-GLM in comparison to the traditional single-neuron GLM. In the first scenario, the external stimulus input 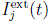 is consistent across trials. Direct connections exist from the source population to the target population. The synaptic weight between each neuron in the source population and each neuron in the target population is random and follows a log normal distribution whose mean is 0.005 and 95% of the weights falls in (0.0023, 0.0109) (underlying normal distribution has a mean of *−*5.29 and standard deviation of 0.4). The self-excitation connections with in each population also follows this distribution. Given that this weight is relatively weak, detecting the connection with a limited number of trials poses a significant challenge for both methods.

In the second scenario, there is no connection between the source and target populations. Instead, the stimulus, which varies from trial to trial, influences both populations. Consequently, the stimulus acts as a confounder in determining the independence of the two populations. We utilized the same piecewise-linear time warping function previously described in the “Modeling trial-to-trial variation in stimulus effects” section to generate a unique stimulus function for each trial. This approach adjusts the peaks of these stimulus functions for both areas to new times, as shown in Figure 4C,D. The correlation between the adjusted peak times in the two areas is 0.5 for the first peak and 0.9 for the second peak, consistent with those observed in the experimental dataset (see the next section). Furthermore, the standard deviations of these shifted peak times are also similar to those observed in the real data.

### 5.4 Experimental dataset and preprocessing

We applied our methodology to the Allen Brain Observatory – Neuropixels Visual Coding dataset [45], which employs high-density extracellular electrophysiology probes, known as Neuropixels, to record spiking activity in various areas of the mouse brain. Recordings were simultaneously obtained from six visual areas using six probes. These areas included the primary visual cortex (V1) and five higher-order visual areas: the lateromedial area (LM), anterolateral area (AL), rostrolateral area (RL), anteromedial area (AM), and posteromedial area (PM). In the experiments, mice were head fixed and passively exposed to a range of visual stimuli including natural movies, flashes, Gabor filters, and drifting gratings. For comprehensive details of the experimental setup, please refer to [2] and [45]. Our study specifically focuses on drifting grating stimuli due to their large number of repeated trials, simplicity, and efficacy in eliciting neural responses. The drifting grating stimuli encompass 40 conditions, comprising combinations of 8 different orientations and 5 temporal frequencies. Each condition was presented 15 times in trials lasting 3 seconds, which included 2 seconds of stimulus exposure followed by 1 second of a gray screen, all arranged in a randomized sequence. For consistency and to minimize potential biases such as p-hacking [58; 59], we selected the same two mouse sessions (757216464 and 798911424) and stimulus conditions as those used by Chen et al. [17]. One session was used for model construction and hyperparameter tuning, while the other was used for validation. The results of both sessions are presented in the Results section. We applied the default spike sorting quality metric filters to the spike trains, which yielded the following neuron counts: In session 757216464, there were 85, 53, 53, 37, 64, and 60 neurons in V1, LM, AL, RL, AM, and PM, respectively. In session 798911424, the counts were 94, 78, 89, 47, 78, and 57. We used 14 and 18 different drifting grating conditions for the two sessions, respectively. The bin size for the spike trains was set at 1 ms. Given that we are analyzing population spike trains, the spike count in each bin is not limited to zero or one. Consequently, 6% of the time bins contain two or more spikes, and 1.5% of the bins contain three or more spikes. This volume of data highlights the computational efficiency of our approach. Because the single-neuron GLM scales quadratically with the number of neurons while pop-GLM maintains constant complexity (relative to population size), the fitting time is reduced from roughly 5 hours to approximately 3 minutes.

To define a neuronal population, we may use all neurons within a visual area or selectively use those neurons that respond to a stimulus and exhibit similar firing patterns. As observed by Chen et al. [17], only a small proportion (20%) of neurons demonstrate strong evoked responses to a stimulus, while the rest show weaker responses or maintain nearly constant firing rates throughout the stimulus periods. We applied the method of Chen et al. [17] to select this subgroup of neurons in our study. A comparison between this selected, strongly evoked population and the full recorded population is presented in S1 Text: the mean firing rates of the two populations are compared in S2 Fig, and the fitted coupling filters are compared in S6 Fig, which shows that the coupling estimates are preserved when all recorded neurons are used.

### 5.5 Excursion algorithm and permutation test

To quantify the discrepancy between two functions, which, in our study, are the coupling filters under stationary and running conditions, we used what has been called an “excursion test” [60]. This method effectively pinpoints regions where the differences between the two functions are most significant. The details of this procedure are outlined in Algorithm 2 and illustrated in Figure S4D. Applying this algorithm to each pair of running and stationary filters yields the observed test statistics. This statistic represents a certain area between the two filters and can be interpreted as the logarithmic geometric mean of spike count increase in the target area per time unit per spike in the source area.

To test the null hypothesis that the coupling filters are identical in both running and stationary trials, we perform a random permutation of the trial types. This permutation test enables us to establish a distribution of the excursion test statistics under the null hypothesis, which allows us to get the statistical significance of the differences observed.

### 5.6 Statistical testing

Several statistical tests are used in this work. In the synthetic-data analyses (Section 3.1), the presence of coupling from the source to the target population is assessed with a likelihood-ratio test (LRT) comparing the full pop-GLM against a nested model in which the source-to-target coupling filter is fixed at zero; for the single-neuron GLMs the per-neuron LRT *p*-values are combined using Fisher’s method, and for reduced-rank regression a permutation test is used. For the experimental data (Section 3.2), differences between the running and stationary coupling filters are quantified with the excursion statistic (Algorithm 2, described above) and assessed with a permutation test over trial labels.

**Algorithm 2:**
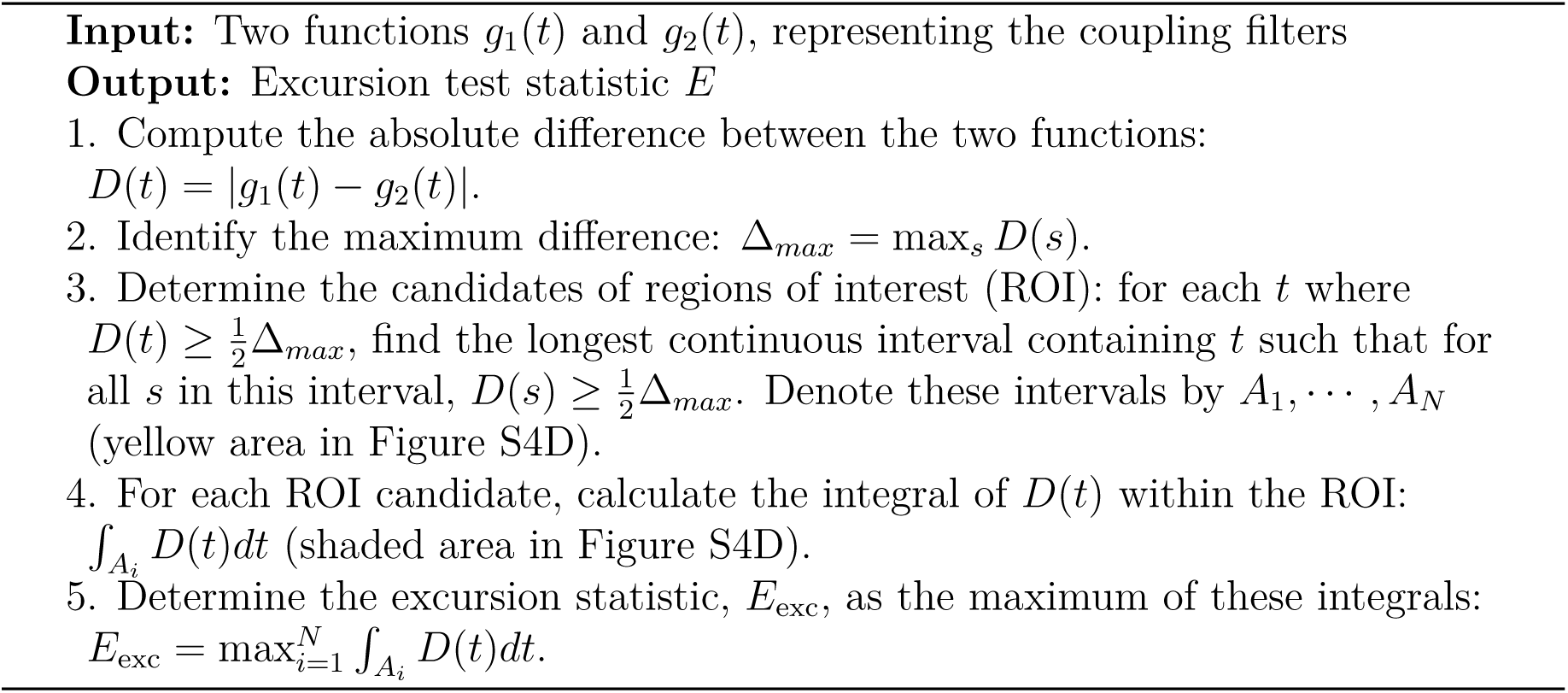
Excursion algorithm.

Because we examine 30 directed pairs of areas, we correct for multiple comparisons. As the 30 tests are not independent, the Bonferroni correction can be conservative [61]; we therefore also compute an exact permutation-based step-down adjustment. In our data the two corrections gave very similar results, consistent with the near-independence of the tests. The exact permutation-based step-down procedure used across the 30 directed area pairs is given as the Testing Procedure algorithm in S1 Text.

## Acknowledgments

Q.X., K.U., and R.E.K. were supported by National Institutes of Health grant R01 MH064537 (to R.E.K.). The funders had no role in study design, data collection and analysis, decision to publish, or preparation of the manuscript. ChatGPT was used by the first author to create an improved initial draft to show to the other authors, who subsequently read through it and edited it thoroughly.

## Data and Code Availability

The Python implementation of the population coupling model (pop-GLM), along with the code used for the simulation and analysis in this paper, is available on GitHub at https://github.com/Qi-Xin/MultiNeuronGLM. The Allen Brain Observatory – Neuropixels Visual Coding dataset used in this study is publicly available via the Allen Institute for Brain Science at https://allensdk.readthedocs.io/en/latest/visual_coding_neuropixels.html.

## Supporting information

**S1 Text.** Supplementary results and methods. Includes: the analytical and empirical demonstration that pooling weakens refractory effects and motivates the nonlinear damping term; model selection for the self-history and damping terms (Table A) and for the running/stationary classification (Table B); the comparison between the selected strongly-evoked population and the full recorded population; and the exact permutation-based step-down procedure used to correct for multiple comparisons.

**S1 Fig.** Explanation of why population spike trains lack refractory effects when using linear self-history filters and require additional nonlinear correction. **A**, Simulate ten independent GLM neurons with a post-spike filter *h* and a constant baseline, then fit a GLM with a self-history filter *g*^ on population spike trains. From left to right: increasing refractory effects. min(*h*) = *−*0.3, *−*1, and *−*3, respectively. The fitted population self-history filter *g*^ is much higher than *h/N_A_* for biologically realistic refractory effects, where a spike would decrease the log firing rate by 2 or 3. **B, C, D**, an example case where an individual neuron does not explode, but a population-level GLM fitted on the population spike train would explode. First, generate spike trains from a population of ten GLM neurons (**B**). There are both post-spike filters for each individual neuron, and a coupling filter from any neuron to any other neuron. Then fit a GLM on the population spike train with two components: inhomogeneous baseline (**C**) and self-history effects (**D**). The fitted population self-history filter is entirely above zero, and we don’t observe any refractory effects. Such strong positive feedback from post-spikes leads the model to explode.

**S2 Fig.** Same as Figure 5, but using all available neurons in each area. Mean firing rates of six populations in six areas, under running and stationary conditions.

**S3 Fig.** Firing rate templates *f_j_*(*t*) from models with only time-warped inhomogeneous baseline 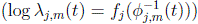. The firing rate template is thinner than the mean firing rate in Fig5.

**S4 Fig. A**, Basis functions for the inhomogeneous baseline template *f_j_*(*t*) consist of 30 B-spline bases. **B**, Basis functions for the cross-area coupling filter *g_i_*_→_*_j_* consist of three Pillow bases. **C**, Basis functions for the population refractory function *f*_damp_Λ consist of four basis functions. They do not start from zero to avoid non-identification issues and are not evenly spaced because there are much fewer data points when Λ is large. **D**, Illustration of excursion statistics. The yellow area shows the region where the difference between the two curves is greater than half the maximum difference, or the region of interest. The excursion statistic is defined as the area of the shaded region.

**S5 Fig.** Simulated PSTH looks similar to experimental PSTH. **A&B**, Experimental PSTH at stationary and running conditions. **C&D**, PSTH of simulated spike trains generated by stationary models and running models.

**S6 Fig. Coupling filters are preserved when all recorded neurons are used instead of the selected populations.** As in Figure 6, but comparing the main-analysis model (pop-GLM, selected populations) with a variant fitted using every recorded neuron in each area. All cross-area coupling filters are shown; each area-pair cell is split into two panels, the *running* condition (left, red) and the *stationary* condition (right, blue). Within each panel pop-GLM (solid line, shaded *±*2*σ* confidence band) is plotted against the *left* axis and the full-neuron variant (dashed line) against the *right* axis, because their amplitude scales differ (right-axis scale *≈* 0.57*×* the left); axis limits are held fixed across all cells. Rows are target areas, columns are source areas (self-coupling omitted). Across the 60 cross-area filters the two agree in shape and timing (correlation of filter integrals *r* = 0.87; of peak latencies *r* = 0.76), while the full-population amplitudes are roughly halved (regression slope 0.50) by dilution from weakly tuned neurons.

**S7 Fig. Coupling filters are essentially unchanged when the nonlinear damping term** *f***_damp_ is replaced by the exact single-neuron self-history.** As in Figure S6 (each area-pair cell split into the *running* condition, left/red, and the *stationary* condition, right/blue; pop-GLM: solid line with shaded *±*2*σ* band; variant: dashed line), but here the variant removes the nonlinear correction term *f*_damp_ and instead accounts for self-history exactly. For each neuron we estimate a self-history filter *h_l_* with a single-neuron GLM using pre-stimulus data (*−*0.5s to 0s, with 0 the stimulus onset), compute the total single-neuron history effect 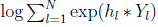 over 0–0.5s, and use it as a fixed offset when fitting the pop-GLM. Because the two models have nearly identical amplitude scales, both share a single axis in each panel. Across the 60 cross-area filters they agree closely in amplitude (correlation of filter integrals *r*= 0.93) and timing (correlation of peak latencies *r* = 0.81). The close agreement indicates that the nonlinear population-level damping term is an adequate approximation to the exact history-dependent effects of individual neurons; the damping term is required for stable simulation but has little influence on the fitted coupling.

**S8 Fig.** Fitted coupling filters using Huk et al.’s method to Allen Institute’s dataset. The coupling filters from V1 to all other areas are the most significant ones and are inhibitory.

**S9 Fig.** Fitted coupling effects’ outputs using Huk et al.’s method. The single-neuron-level model suggests that V1’s inhibition to other areas dominates the firing rates. The two firing rate peaks result from V1’s low activity tens of milliseconds ago. These fitted results do not seem reasonable.

**S10 Fig. A**, Trial-averaged Local Field Potential (LFP) post-stimulus onset under both stationary and running conditions. The LFP is averaged across multiple channels within an area. Left: V1. Right: LM. Under stationary conditions, two peaks are observed, whereas the second peak is less prominent when running, aligning with the spiking intensity observed in Figure 5. **B**, Trial-averaged Granger causality test statistics from V1 channels to LM channels. Left: stationary conditions. Right: running conditions. The Granger causality test statistics reveal a stronger connection from V1 to LM than from LM to V1, and these statistics notably diminish during running. The order of the Granger causality test is five, determined by AIC, and we ensured the significance in the tests is not due to nonstationarity by detrending and visualization.

**S11 Fig.** Fitted coupling filter from V1 to LM for the second validation session. We directly applied the analysis from the first session without altering any hyperparameters. The significant connection change from V1 to LM, observed in the first mouse, is also substantiated in the second mouse.

**S12 Fig. A**, Simulated firing rates of six areas using models with and without the nonlinear correction term *f*_damp_. **B**, Fitted *f*_damp_ in the six areas under stationary and running conditions.

**S13 Fig. Coupling and trial-to-trial baseline variation both present (synthetic Scenario 3).** EIF populations were simulated with a nonzero feed-forward coupling *and* across-trial jitter in the timing of the two response peaks (all settings as in the Sensitivity scenario); a representative simulation of 100 trials is shown. **A**, pop-GLM coupling filter fitted with time warping (black) and without it (red), each with its 95% confidence interval (shaded). Without time warping the filter is inflated—its confidence interval lies higher—because the unmodelled trial-to-trial variation is misattributed to coupling. **B**, single-neuron GLM coupling filters (grey; a random subset of the 100 fitted filters) and their mean (black); these are far noisier than the pop-GLM estimate and are also inflated, because an individual neuron carries too little information to estimate per-trial shifts and so cannot implement time warping.

**S14 Fig.** Fitted GLM outputs of each component, averaged across trials (95% CI). Only coupling effects and time-warped inhomogeneous baselines are shown. Red: running. Blue: stationary. Excursion tests were performed on the two curves in each panel, but the null distribution is not shown here. Yellow areas indicate the selected regions found by excursion tests with *p*-values below 0.05. For example, the first row shows the response variable as V1 population spike trains, with coupling effects from all six areas and an inhomogeneous baseline (interpreted as input from the thalamus and other unobserved sources).

