## Supplementary for "A Population Coupling Model Identifies Reduced Propagation from V1 to Higher Visual Areas During Locomotion": s1 text.pdf

### Supplementary materials

The supplementary material is organized into *Supplementary Results* and *Supplementary Figures and Tables*, ordered to follow the logic of the main text: model design, synthetic validation, and biological application.

### Supplementary Results

#### Model design: why population spike trains require a nonlinear damping term

##### Lack of refractory effects in linear self-history filters of population spike trains

In this section, we show that the refractory effect, a transient period after a spike during which the neuron's likelihood to fire again is reduced, can become less pronounced at the population level after pooling individual spike trains. This discrepancy can have significant consequences when modeling with GLMs. Specifically, a population-level model may exhibit instability even if individual neurons are stable.

Consider a population consisting of $N_A$ GLM neurons. Each neuron in the population only receives self-history effects from itself. For simplicity, we will assume that all neurons share an identical self-history filter $h$ and a constant baseline $\beta_0$.

For each neuron, its behavior can be described as:

$$\log \lambda_i = h * Y_i + \beta_0, \tag{1}$$

where $\lambda_i$ is the firing rate of the i-th neuron, and $Y_i$ represents its binary spike train.

Let $Y = \sum_{i=1}^{N_A} Y_i$ denote the population spike train. The population firing rate is given by

$$\lambda = \sum_{i}^{N_A} \lambda_i. \tag{2}$$

We can derive the firing rate of the population spike train as:

$$\log \lambda = \log \left( \sum_{i=1}^{N_A} \lambda_i \right) \tag{3}$$

$$= \beta_0 + \log \left( \sum_{i}^{N_A} \exp(h * Y_i) \right). \tag{4}$$

Due to the convexity of exponential function, we have the following inequation

$$\sum_{i}^{N_A} \exp(h * Y_i) \geq N_A \cdot \exp(h * Y/N_A). \tag{5}$$

Thus,

$$\begin{aligned} \log \lambda &\geq \log(N_A) + \beta_0 + h * Y/N_A \qquad (6) \\ &= \tilde{\beta}_0 + Y * \tilde{h}, \qquad (7) \end{aligned}$$

where $\tilde{\beta}_0 = \log(N_A) + \beta_0$ and $\tilde{h} = h/N_A$ represent the linearly scaled baseline and post-spike filter.

This inequality implies that the true log firing rate of the population, $\log \lambda$, is greater than the prediction provided by the linearly scaled parameters $\tilde{\beta}_0$ and $\tilde{h}$. Therefore, if we fit a population GLM to the data to obtain estimated parameters $\hat{\beta}_0$ and $\hat{h}$, these estimates must compensate for this underestimation. Consequently, compared to the theoretically scaled parameters $\tilde{\beta}_0$ and $\tilde{h}$, at least one of the following assertions regarding the fitted estimates must hold true:

- The estimated baseline is higher than the linearly scaled prediction ($\hat{\beta}_0 > \tilde{\beta}_0$).
- The estimated population self-history filter $\hat{h}$, which is typically negative during the refractory period, is weaker (less negative) than the scaled individual filter $\tilde{h}$.

In our simulation, we consistently observed that the fitted population self-history filter, denoted by $\hat{h}$, is substantially weaker than $\tilde{h}$ when the post-spike filters of individual neurons emulate biologically authentic refractory effects (Figure S1A). Consequently, the overall refractory effects in the population-level GLMs, given by $\hat{h} * \tilde{Y}$, are much weaker than $\tilde{h} * \tilde{Y} \approx h * Y_i$, which represents the refractory effects in a single neuron. This suggests that there tends to be a lack of refractory effects if we just use a linear self-history filter to capture refractory effects in population spike trains.

Next, we highlight a scenario where an individual neuron remains stable, but the model tailored to the population spike trains exhibits instability due to this lack of refractory effects. In this model, we introduce excitatory coupling effects within the population of $N_A$ GLM neurons. Similarly, we assume that every neuron receives coupling filters from all other neurons in the group and all coupling filters are consistent, represented by $c$. Figure S1 B shows the generated spike trains, where the ground truth single-neuron GLM never explode. However, if we fit population-level GLM with a linear self-history filter and inhomogeneous baseline to the population spike train, it becomes unstable, as the fitted self-history filter does not reflect strong refractory effects and the self-history filter is dominated by excitatory coupling effects (Figure S1D). The fact that linear self-history filters at population level fail to capture enough refractory effects explains why the nonlinear correction term in our pop-GLM is very significant and prevents explosion.

#### Model Selection for Self-History Effects

This section provides a detailed exploration of the model selection process, specifically focusing on addressing the explosion issue (Figure 1C). These explorations are summarized in Table A, where various model configurations are compared based on their Bayesian Information Criterion (BIC) improvements and stability. The BIC is defined as $\mathrm{BIC} = -2\log\hat{L} + k\log n$, where $\hat{L}$ is the maximized likelihood, $k$ the number of free parameters, and $n$ the number of observations; a lower BIC indicates a better penalized fit, and the values reported here and in the following table are improvements in BIC relative to a baseline model.

Initially, we identified that both cross-area coupling and self-history effects are necessary and significantly improve the model's fit. However, as indicated in the main text, adding self-history effects led to instability. To address this, we explored various modifications to the model.

One approach was the inclusion of additional terms to the log firing rate. We introduced a new covariate, $\Lambda(t) = \frac{1}{\tau}\sum_{k=1}^{n}\exp(-(t-t_k^*)/\tau)$, capturing the empirical spike density over a short history window. Variations of this term were tested, including $\Lambda$ itself, its quadratic form $\Lambda^2$, and a nonparametric term $f_{\text{damp}}(\Lambda)$. Among these, the nonparametric term $f_{\text{damp}}(\Lambda)$ stood out for its robust ability to prevent explosion and significantly improve the fit, as detailed in the main text and shown in Table A.

Another pathway we explored was adding the dependence between the empirical spike density $\Lambda(t)$ and self-history filters, rather than directly adding a term to the log firing rate. Specifically, we modified the fixed self-history filter $\mathbf{g}$ to a varying version $\widetilde{\mathbf{g}(\Lambda)} = \mathbf{g_0} + \Lambda\cdot\mathbf{g_1}$, which depends on $\Lambda$, allowing for more flexible adaptation to the spike train's history. In this way, the total self-history effects becomes:

$$
\begin{aligned}
h(t|H_t) &= \mathbf{Y} * \tilde{\mathbf{g}} &(8)\\
&= \mathbf{Y} * (\mathbf{g_0} + \Lambda\cdot\mathbf{g_1}) &(9)\\
&= \mathbf{Y} * \mathbf{g_0} + \Lambda\cdot\mathbf{Y} * \mathbf{g_1} &(10)\\
&= \sum_{k=1}^{n} g_0(t-t_k^*) + \Lambda\cdot\sum_{k=1}^{n} g_1(t-t_k^*). &(11)
\end{aligned}
$$

We also considered second-order interactions by substituting $\mathbf{g}$ with a more complex formulation of $\tilde{\mathbf{g}}(\Lambda) = \mathbf{g_0}+\Lambda\cdot\mathbf{g_1}+\Lambda^2\cdot\mathbf{g_2}$. Although incorporating first-order dependency improved the fit and stabilized the model, it did not surpass the enhancements brought by the nonparametric term added to the log firing rate. Therefore, for its robustness, adaptability, and simplicity, we ultimately chose the nonparametric term $f_{\text{damp}}(\Lambda)$ as our final model.

| Model components | | | | BIC improvement | Stability |
|---|---|---|---|---|---|
| Baseline | Coupling | Self-history | Correction term | | |
| ✗ | | | | 0 | Yes |
| ✗ | ✗ | | | 4362.17 | No |
| ✗ | ✗ | ✗ | | 5941.72 | No |
| ✗ | ✗ | ✗ (with first-order dependence) | | 8183.60 | Yes |
| ✗ | ✗ | ✗ (with second-order dependence) | | 7682.57 | No |
| ✗ | ✗ | ✗ | ✗ (linear) | 6383.48 | No |
| ✗ | ✗ | ✗ | ✗ (quadratic) | 8094.84 | Yes |
| **✗** | **✗** | **✗** | **✗ (nonparametric)** | **8929.0** | **Yes** |

**Table A**: Model selection for the self-history and correction terms. Candidate models for the V1 (probe C) population spike train, compared by Bayesian Information Criterion (BIC) and by stability under simulation. A mark in a column indicates that the component is included in that model. All models contain the time-warped inhomogeneous baseline together with the trial-wise gain constant ("Baseline"); "Coupling" adds the coupling filters convolved with the population spike trains of the other five areas, and "Self-history" adds the post-spike filter convolved with the population's own spike train. The rows marked *with first-order* and *with second-order dependence* replace the fixed self-history filter $\mathbf{g}$ by a $\Lambda$-dependent filter, $\tilde{\mathbf{g}}(\Lambda) = \mathbf{g_0} + \Lambda \cdot \mathbf{g_1}$ and $\tilde{\mathbf{g}}(\Lambda) = \mathbf{g_0} + \Lambda \cdot \mathbf{g_1} + \Lambda^2 \cdot \mathbf{g_2}$ respectively. The three "Correction term" rows instead add a further predictor to the right-hand side of the log firing rate: $\Lambda$ itself (linear), $\Lambda^2$ (quadratic), or the nonparametric function $f_{\mathrm{damp}}(\Lambda)$ used in pop-GLM. Running and stationary trials were fitted as separate models and their BIC values summed. "BIC improvement" is the reduction in BIC relative to the baseline-only model in the first row, so that larger values indicate a better penalized fit. Stability was assessed by simulating spike trains from each fitted model; models that were either fragile (occasionally explode) or divergent (always explode) are labelled "No". The nonparametric correction term $f_{\mathrm{damp}}(\Lambda)$ (bold, last row) yields both the largest BIC improvement and a stable model, and is the specification used throughout the paper.

#### Selected versus full recorded population

Figures 5 and S2 compare the mean firing rate per neuron in each area when including only the identified subpopulation versus all available neurons. Although the overall shapes of the firing-rate functions are largely unchanged, the amplitudes decrease to about a quarter of their original values when all neurons are included. The inclusion of non-relevant neurons essentially introduces additional Poisson noise, lowering the per-neuron firing rate, and also dilutes the observed coupling strength in single-neuron analyses. Figure S6 shows the fitted coupling filters with all neurons included: the shapes are similar to those using just the subpopulation (Figure 6), but the amplitudes are reduced. A quantitative comparison across all 60 cross-area coupling filters confirms that the results are preserved under the full-population choice: the filters agree in shape and timing (correlation of filter integrals $r = 0.87$; correlation of peak latencies $r = 0.76$), while their amplitude is roughly halved (the variant is on average 0.50 times the main model), as expected from dilution by weakly tuned neurons (Supplementary Figure S6). All model variants reproduce the reduction of V1-to-LM coupling during locomotion.

#### Model selection for running versus stationary classification

We performed model selection for the first mouse to determine the optimal incorporation of running speed effects into our model. Details are given in Table B, but note that the models did not include coupling effects (the variables of interest). We discovered that categorizing trials into "running" and "stationary" based on average speed was the most effective, which is typical in the literature [1; 2; 3; 4]. Specifically, a trial is labeled as "running" if the average speed exceeds 1 cm/s, while those with lower speeds are classified as "stationary". Adopting stricter criteria as in [4] to identify these type of trials resulted in similar outcomes, but with reduced sample sizes. Simplifying the speed effects into just two trial types also allows us to fit separate models only to two conditions, which makes both the fitting process and subsequent inference easier.

### Supplementary algorithm

#### Testing procedure for the multiple-comparisons correction

The exact permutation-based step-down procedure used to correct for multiple comparisons across the 30 directed area pairs—referred to from the main-text Statistical testing section—is detailed in the Testing Procedure algorithm below.

| Model name and equation | BIC improvement comparing to baseline |
|---|---|
| Baseline without considering speed:<br>$\log \lambda_t = f_t$ | 0 |
| Linear term of instantaneous speed:<br>$\log \lambda_t = f_t + \beta \cdot s_t$ | 75.09 |
| Binary term of instantaneous binary state:<br>$\log \lambda_t = f_t + \beta \cdot I(s_t > v_{th})$ | 6.93 |
| Two-way coupling of instantaneous speed:<br>$\log \lambda_t = f_t + \mathbf{k}_{speed} \cdot \mathbf{s}_{(t-l):(t+l)}$ | 109.12 |
| Time-dependent term of instantaneous speed:<br>$\log \lambda_t = f_t + \beta_t \cdot s_t$ | 555.69 |
| Time-dependent term of instantaneous binary state:<br>$\log \lambda_t = f_t + \beta_t \cdot I(s_t > v_{th})$ | 914.43 |
| **Time-dependent term of trial-wise binary state:**<br>$\boldsymbol{\log \lambda_t = f_t + \beta_t \cdot I(\bar{s} > v_{th})}$ | **1454.99** |

**Table B**: Model selection with regard to speed. In our model selection process concerning speed, we fit models incorporating the effects of speed in various ways and compare their Bayesian Information Criterion (BIC). The model with the trial-wise binary state provides the best BIC, which is highlighted in bold. According to this model, a trial is classified as a running trial if its average speed exceeds 1 cm/s. Conversely, trials with an average speed below this threshold are classified as stationary. In our notation, $\lambda_t$ denotes the firing rate at the t-th time bin, and $f_t$ is the time-varying baseline. The term $s_t$ represents the recorded running speed at time t, and $\beta$ is the fitted coefficient. For defining the binary state, $I$ is the indicator function, and $v_{th}$ is the speed threshold used to classify states as either stationary or running. The vector $\mathbf{k}_{speed}$ serves as the filter associated with the recent past and future running speeds. $\beta_t$ is the non-stationary coefficient related to instantaneous speed or state. In all models listed in the table, we have also considered the optimal lead-lag time between the firing rate $\lambda_t$ and running speed $s_t$, as well as the best smoothing parameter for running speed $s_t$, determined through cross-validation. $v_{th}$ is another hyperparameter, which was optimized using cross-validation. In practice, a $v_{th}$ value of 1 provided the best fit, resulting in an equal distribution of running and stationary trials.

**Algorithm 1:** Testing Procedure

**Input:** Test statistics for 30 cases under 82600 permutations $T_{i,n}$ $(i = 1, 2, ..., 30;\ n = 1, 2, ..., 82600)$; observed test statistics for 30 cases $T_i^{obs}$

**Output:** Corrected $p$-values $p'_{(1)}$, $p'_{(2)}$, ... for the most significant cases

1. Calculate the nominal $p$-value for each case. Identify the smallest nominal $p$-value, denoted by $p_{(1)}$
2. For each permutation $n = 1, 2, \ldots, 82600$, compute the nominal $p$-values for the set of test statistics $T_{:,n}$. Record the frequency at which the smallest $p$-value is less than $p_{(1)}$, which is denoted as the corrected $p$-value $p'_{(1)}$.
3. Exclude the case associated with $p_{(1)}$, leaving 29 cases. Repeat step 2, using $p_{(2)}$ in place of $p_{(1)}$, to determine $p'_{(2)}$.
4. Continue as in step 3 for the i-th smallest $p$-value $p_{(i)}$, until the corrected $p$-value $p'_{(i)}$ exceeds 0.05.
